# Historical Contingency Limits Adaptive Diversification in a Spatially Structured Environment

**DOI:** 10.1101/2025.07.27.667043

**Authors:** Gillian Patton, John C. Meraz, Michelle Yin, Sarah B. Worthan, Sara Williams, Megan G. Behringer

**Affiliations:** Department of Biological Sciences, Vanderbilt University, Nashville, Tennessee, USA; Biodesign Center for Mechanisms of Evolution, Arizona State University, Tempe, Arizona, USA; Department of Pathology, Microbiology, and Immunology, Vanderbilt University Medical Center, Nashville, Tennessee, USA; Evolutionary Studies Initiative, Vanderbilt University, Nashville, Tennessee, USA

**Keywords:** range expansion, experimental evolution, historical contingency, fitness landscape, evolutionary trap

## Abstract

Understanding how genotype-by-environment (GxE) interactions influence evolutionary trajectories and contribute to historical contingency is key to predicting evolution. In spatially structured, heterogeneous environments, populations often diversify into ecotypes, resulting in niche specialization. However, the ability to specialize depends not only on ecological opportunity but also on whether genetic variation permits access to novel niches, as genetic disruptions may inhibit adaptation unless alternative trajectories or compensatory mutations are subsequently accessible. Previously, we demonstrated that *Escherichia coli* populations rapidly diversify into two co-existing ecotypes in a nutrient-rich, spatially structured environment. The adaptation of both ecotypes results in benefits only perceived in the spatially structured culture tube environment. Here, diversification is initiated by first-step mutations associated with the overexpression of genes encoding the type 1 fimbria, the major attachment pilus involved in biofilm development, enabling range expansion and allowing *E. coli* to occupy the surface-air interface of the culture tube.

To investigate how first-step mutations shape evolutionary trajectories, we experimentally evolved wild-type and fimbrial-deficient (Δ*fimA*) *E. coli* for 91 days in both structured (tube) and unstructured (flask) environments. While a *fimA* deletion initially confers a fitness benefit by avoiding the cost of insufficient biofilm formation, it ultimately prevents range expansion in structured environments and is not compensated by expression of cryptic fimbriae by the end of experimental evolution. As a result, Δ*fimA* populations show constrained adaptation in tubes compared to wild-type. Alternatively, both genotypes perform similarly in flasks, where biofilm formation is not advantageous and whole population sequencing reveals that flask-evolved populations similar early mutational trajectories. Our results highlight the ruggedness of the adaptive landscape in structured environments and show how an initially beneficial mutation can trap a lineage on a local fitness peak, underscoring the importance of G×E interactions and early mutational events in shaping the predictability and contingency of evolutionary outcomes.

## Introduction

When faced with a novel environment, organisms must adapt in order to thrive. However, populations with similar genetic backgrounds may follow different adaptive trajectories due to the influence of chance and evolutionary history (1–3). Here, the term “chance” includes novel mutations and genetic drift, which stochastically shape genetic variation (4–6). History is the genetic and developmental background of the evolving population, which can limit the accessibility of different adaptive trajectories (7–10). Because an early beneficial mutation could render an otherwise more optimal adaptive trajectory inaccessible and change the direction of evolution, a population’s adaptability, or capacity to adapt to a specific environmental condition, is highly contingent upon history and chance (11–15).

The repeatability of how natural selection mediates history and chance while navigating the adaptive landscape can be investigated through the experimental evolution of microbial populations (16). Microbes are an ideal system as their large population sizes and quick generation times allow for the assessment of evolutionary processes over relatively short timescales. A common observation in experimental evolution studies is genetic parallelism between populations that evolved in similar environments, demonstrating that shared selective pressures result in the evolution of genetic similarities (17–20). However, even under extreme selection pressures, such as sub-lethal stress, evolution is not 100% repeatable. One reason is genotypic redundancy, where multiple mutations can produce the same phenotype (21–23). Genetic incompatibilities can further limit parallelism as two independently beneficial mutations can exhibit sign epistasis, conferring reduced fitness when combined (24). Such incompatibilities can constrain populations to very different evolutionary trajectories, which can curtail adaptation if a population becomes stuck on a locally adaptive peak (25, 26). In a complex environment containing multiple spatial niches, a population’s inability to expand its range and inhabit a new niche may constrain its ability to evolve increased fitness (27, 28). Thus, if an early beneficial mutation ultimately limits range expansion, or the ability of a species to expand its distribution and colonize new niches, it may lead to an evolutionary dead end (29–31). To examine how genotype and environment interact to shape adaptability, we will assess how a single mutation can affect adaptation to different culture environments using biofilm formation in *Escherichia coli* as a model.

Biofilms are collaborative interactions of bacterial cells that confer tolerance to stress, antibiotics, and host immunological defenses (32). *Escherichia coli*, like many other species of bacteria, forms biofilms in environments where collaboration may be necessary to take advantage of a particular niche or where protective measures are required (33–35). In addition to defense, biofilms confer many other benefits, including greater access to oxygen at the surface-air interface of a liquid, limitation of competition, and facilitation of cooperation (36–40). In laboratory environments, *E. coli* can form pellicle biofilms at the surface-air interface through adhesion with cell surface appendages called fimbriae (33, 41, 42). The *E. coli* genome contains 14 different operons that encode adhesins, including type 1 fimbriae (*fim*), curli fimbriae (*csg*), antigen 43 (*flu*), and the *E. coli* common pilus (*ecp*) (43, 44). Although the genes for the other fimbrial adhesins are intact and functional when genetically engineered to be expressed, *E. coli* K-12 MG1655 only expresses the *fim* and *flu* operons under standard laboratory conditions (45). Biofilm formation via the *fim* operon is a phase-variable phenotype whereby two site-specific recombinases, FimB and FimE, control expression by inverting the *fim* promoter (*fimS*) into the “on” or “off” orientation (46, 47). *fimA* is the most highly expressed of the seven genes in the *fim* operon and is essential to type 1 fimbriae formation (48). As a typical type 1 fimbria can contain ∼1,000 subunits of FimA, and a fimbriated cell can express 200 - 500 fimbriae on its surface, expression of the *fim* operon and forming a biofilm represents a substantial energetic commitment, especially when *E. coli* K-12 MG1655 cells are generally weak biofilm formers in standard laboratory conditions and are unable to fully realize the spatial and oxygenation benefits of biofilm formation (48, 49).

Despite the steep energetic costs, we previously observed that mutations causing increased biofilm-forming activity through the overexpression of the *fim* operon are a common early adaptive step during the experimental evolution of *Escherichia coli* populations in culture tubes (50–52). In these populations, biofilm formation precedes diversification as *E. coli* evolves to expand its range within the culture tube by colonizing both the highly aerobic surface-air interface and the less aerobic planktonic/bottom-dwelling niche (50). As such, *E. coli* biofilm formation represents a useful model for studying adaptability via eliminating access to a first-step mutation. By deleting *fimA* and thus preventing biofilm formation via the *fim* operon, we can then repeat evolution and survey the adaptive landscape to identify alternative evolutionary trajectories and how these trajectories translate into increased fitness (11, 22, 53).

Moreover, because the fitness of a particular genetic background is dependent on the environment (54–56), differences between culture vessels, such as their shape and aeration capabilities, can significantly alter *E. coli*’s physiology and, by extension, its evolution (57). A well-shaken culture flask is essentially a non-structured environment where the medium is more oxygenated and homogenized, while a culture tube is a much more structured environment, where the presence of an oxygen gradient can give rise to the creation of additional distinct niches, simulating some of the complexity of natural environments. If biofilm formation is essential to adaptation in culture tubes, we expect the *E. coli* populations would evolve to activate one of the other 11 cryptic fimbriae operons (45). Alternatively, if *E. coli* populations do not activate cryptic fimbriae, either biofilm formation is not essential to adaptation in culture tubes or mutations that enable cryptic fimbriae are not easily accessible. As biofilm formation provides limited benefits in culture flasks due the homogeneity of the environment, we’d expect minimal differences in adaptability based on a population’s initial biofilm-forming potential (58).

In this study, we investigate how interactions between genetic background and environment influence adaptability by evolving *E. coli* populations with two different initial genotypes, WT (biofilm-enabled) and Δ*fimA* (biofilm-inhibited), in two distinct environments, culture flasks (no spatial structure) and culture tubes (spatially-structured), for 91 days (13 weeks, 215 generations) (**Figure 1**). Throughout this experiment, we measured changes in culture density and biofilm production. After 91 days of evolution, we assessed changes in fitness suggesting differences in local adaptation while whole-population metagenomic sequencing reveals genotype-by-environment effects on molecular evolution.

**Figure 1:**
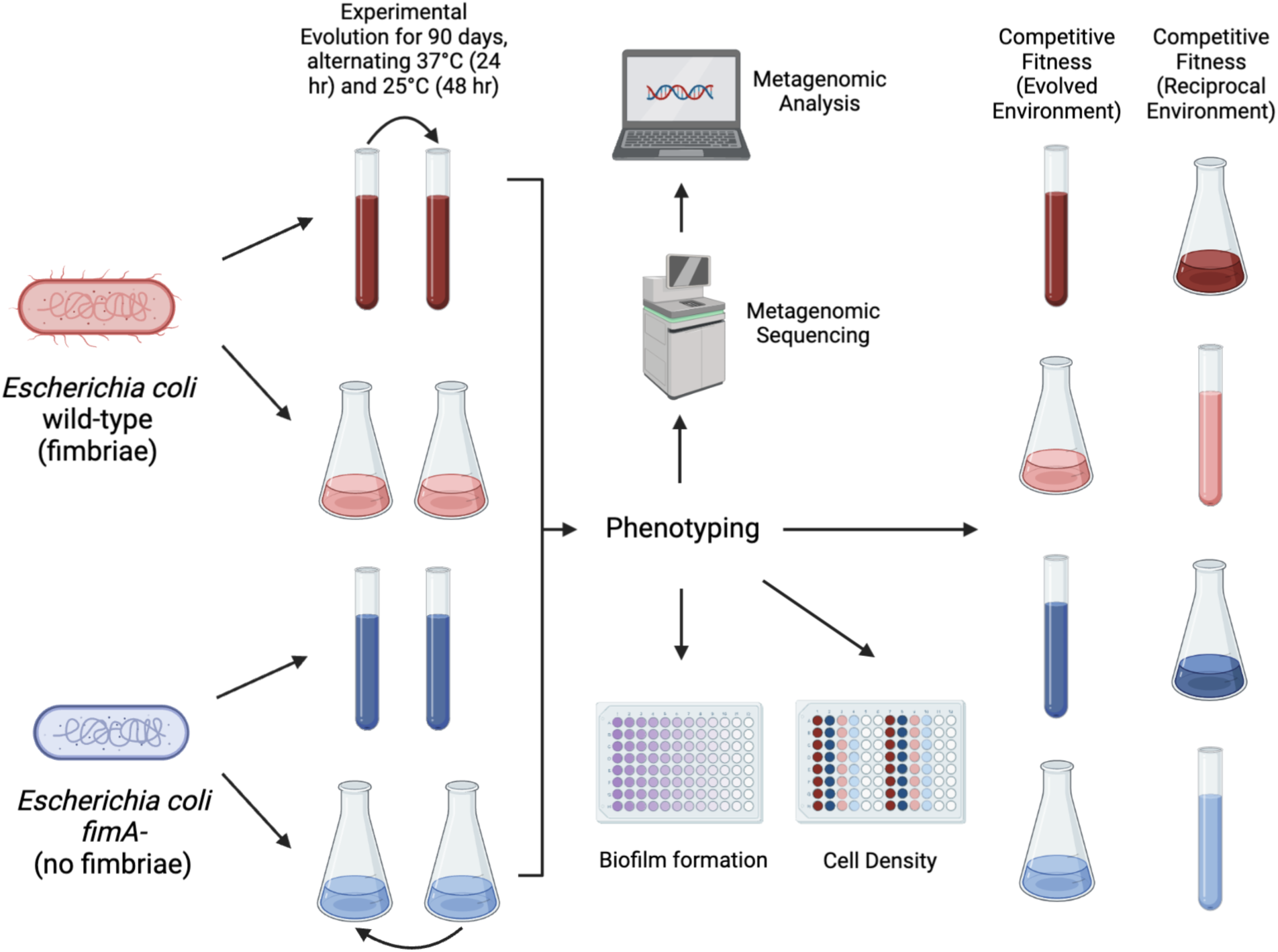
Illustration of experimental methods. In *Escherichia coli* K-12 strain MG1655, we experimentally evolved a total of 24 parallel populations split across 4 different treatments: WT-tube WT-flask, *ΔfimA*-tube, and *ΔfimA*-flask. Transfers were conducted alternating 48h of growth at 25°C and 24h of growth at 37°C, shaking at 180 rpm. Each transfer consisted of a 1:10 dilution where 1 mL of well-mixed evolving culture is inoculated into 9 mL of fresh LB broth; cell density was measured weekly over the course of evolution. After 91 days of experimental evolution (13 weeks), the evolved populations were compared to their ancestors via biofilm formation assays, competitive fitness assays, and metagenomic analysis.

## Results

### Environmental conditions have a distinct effect on biofilm formation and culture density

As increased biofilm formation via overexpression of *fimA* is often the first phenotype observed when *E. coli* evolves in a structured culture tube environment (50–52), we replayed evolution to determine how adaptation is affected when evolving populations are unable to form biofilms. We deleted the major subunit of type 1 fimbriae (*ΔfimA*) from the WT experimental ancestor strain to create a biofilm-deficient experimental ancestor and assessed the baseline biofilm production of both genetic backgrounds in a 96-well plate (59). Baseline assessment of biofilm production confirmed that the *ΔfimA* mutant did not produce biofilm in either culture condition, while consistent with previous reports, growth for 48h at 25°C induces biofilm formation in the WT ancestor (60)(ANOVA with Tukey’s HSD, WT: *P*_37°C vs. 25°C_ =4.82 x 10^-10^; **Figure 2A**, **Figure S1**).

**Figure 2:**
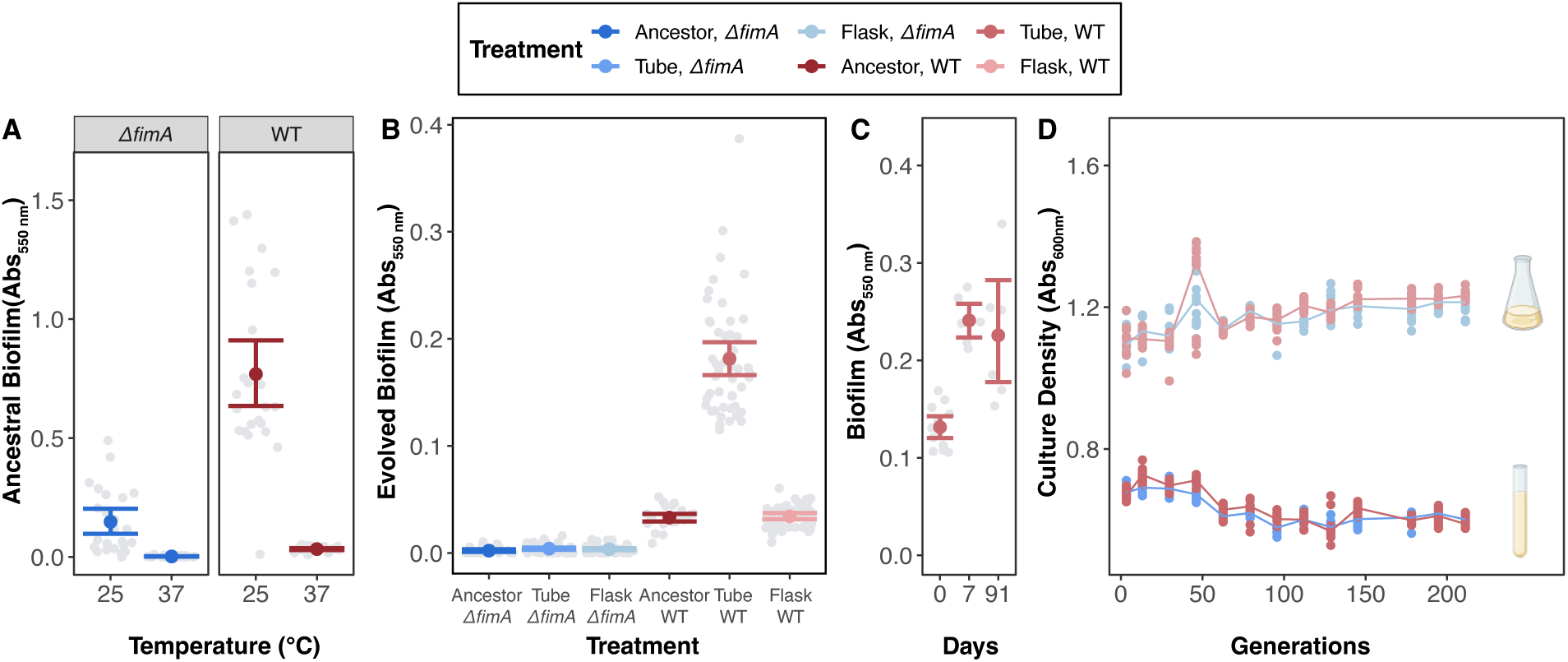
Biofilm production increases only in WT populations experimentally evolved in the tube environment. Biofilm production was assessed following static growth in a 96-well plate. (A) Ancestral capacity for biofilm production after 48h at 25°C and 24h at 37°C. (B) Biofilm production after 91 days of experimental evolution, and C) biofilm production of WT-tube populations measured after 0, 7, and 91 days of experimental evolution. D) Pre-transfer density of cultures assessed every 7 days throughout the evolution experiment. For panels A-C, light grey circles represent measurements of each population (8 replicates). Colored circles indicate the mean for each treatment and error bars represent 95% confidence intervals of the mean. For panels B-D, biofilm or culture density was assessed after 24h at 37°C (48h at 25°C is presented in **Figures S1-2**). For panel D, colored circles represent the cell density measurements of each population and colored lines connect the means of each timepoint to visualize trends.

We then initiated 24 experimental populations from the WT and *ΔfimA E. coli* strains. Each population was cultivated via serial-transferred batch culture over 91 days (215 generations) in either a heterogeneous culture tube environment or a more homogeneous culture flask environment (four genetic background/culture condition treatments; six populations each) and after 91 days of experimental evolution, we re-assessed the populations’ ability to form biofilms. Here, we found that all *ΔfimA* populations remained unable to produce biofilms while WT-flask populations lost their ancestral ability to produce biofilms during 48h growth at 25°C (**Figure 2B**, **Figure S1**). In contrast, the WT-tube populations gained the ability to form biofilms during 24h growth at 37°C (ANOVA with Tukey’s HSD, WT-tube*: P* = 1.78 x 10^-11^), and this increase in biofilm formation could be observed as early as the day 7 timepoint (ANOVA with Tukey’s HSD, *P*_Day 0 vs. 7_ = 4.83 x 10^-5^; *P*_Day 7 vs. 91_ = 0.78; **Figure 2C**). Given this rapid change in phenotype, we expect the initial increase in biofilm is due to retention of the phase variable expression of the *fim* operon following the first iteration of 48h growth at 25°C.

As any evolved increases in culture density could skew the ability to detect differences in biofilm formation, we tracked the evolution of culture density weekly throughout the experiment. Although both WT and *ΔfimA* populations evolved changes in culture density, these changes were restricted to the 24h/37°C transfers and are unlikely to be a contributing factor to the evolved differences in biofilm formation as populations that evolved in tubes are consistently less dense than populations that evolved in flasks (ANOVA with Tukey’s HSD, *P*_tubes vs. flasks_ = 4.79 x 10^-10^, **Figure 2D, Figure S2**). This reduced culture density in tubes is consistent with slower growth due to heterogeneous oxygen and resource conditions in culture tubes (57).

### Inability to form biofilms has varying effects on evolved fitness based on environment

Since we observed a notable increase in biofilm formation from WT-tube populations as early as day 7 that was retained throughout experimental evolution, we suspect that an ability to increase biofilm production is likely an important trait for adaptation in the culture tube environment. To determine how the ability to produce biofilm influences competitive fitness, we co-cultured the evolved populations against their experimental ancestor in their respective evolution conditions (57). Overall, the greatest changes in fitness are observed in the 24h/37°C culture conditions where all evolved populations exhibit increased fitness relative to their ancestor (**Figure 3A**). Notably, the mean fitness for *ΔfimA-*tube populations is only 1.04 which is within the range of error of the *ΔfimA*-ancestor and quite modest compared to all other evolutionary treatments whose fitness range from 1.29 - 1.40 in 24h/37°C conditions. As marginal changes in fitness are observed for *ΔfimA* populations cultivated in tube but not flasks, this suggests that an inability to produce biofilms may only hinder adaptation in the tube environment. Alternatively, fitness was modest across all evolved populations in the 48h/25°C culture environment with only WT-flask populations exhibiting significantly greater fitness than its ancestor (*P*_WT-flask_ = 0.54, **Figure S3**). This bias in evolved changes contributing to greater fitness gains in the 24h/37°C conditions, instead of both conditions, highlights the specificity of adaptation and suggests that the greatest advantage can be obtained through evolved changes that confer benefits specific to the 24h/37°C condition. As such, we focused our following investigations of fitness on these conditions.

**Figure 3:**
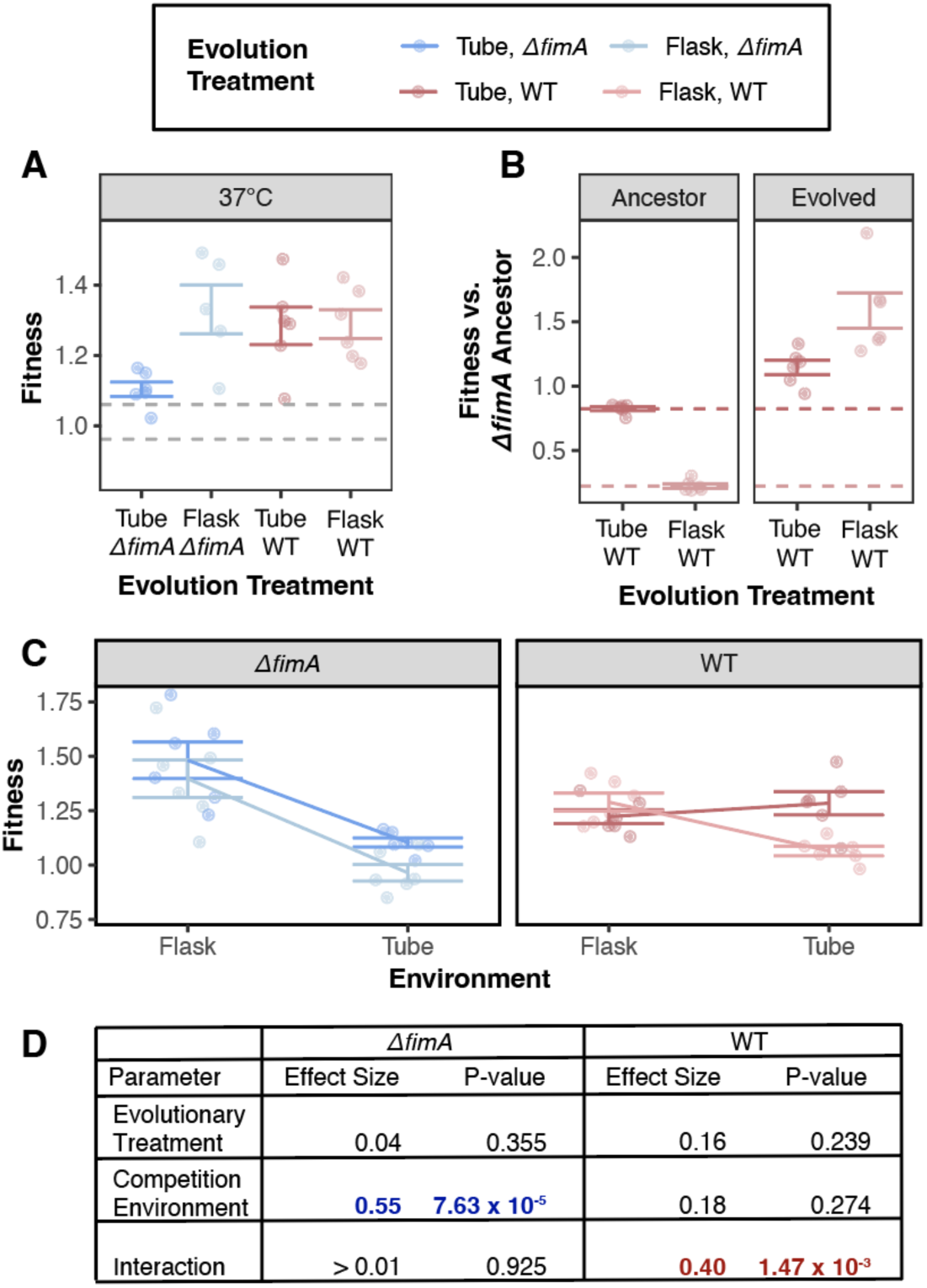
Loss of *fimA* creates an advantage in homogeneous environments and a disadvantage in heterogeneous environments. Fitness is assessed by competitive co-culture in evolved and reciprocal environments. (A) Fitness of each population in their evolved environment after 91 days relative to the ancestor strain of the same genetic background (e.g. evolved *ΔfimA* tube with *ΔfimA* ancestor). (B) Fitness of ancestral (left) and evolved (right) wild-type populations relative to the *ΔfimA* ancestor assessed by competitive co-culture. (C) Fitness of each population in their evolved or reciprocal environment relative to the ancestor strain of the same genetic background. Lines illustrate fitness relationships by connecting the mean fitness for each evolved genotype-environment combination. For panels A-C, fitness assays were conducted at 37°C (25°C fitness assays are presented in **Figure S3**); colored circles indicate fitness values for each population, and error bars represent 95% confidence intervals of the mean. (D) Effect size of evolutionary treatment and competitive environment for each genetic background cultured in their evolved or reciprocal environment determined via multiple linear regression. Effect size is calculated as partial eta^2^ and significant parameters are highlighted in blue (*ΔfimA)* or red (WT).

Given the modest magnitude of fitness evolution exhibited by *ΔfimA-*tube populations, we considered the possibility that the *ΔfimA* experimental ancestor may have started the evolution experiment closer to the fitness optimum than the WT experimental ancestor. Here, a more limited fitness increase would be expected due to diminishing returns epistasis compared to the contrasting extreme of significant fitness gains by WT-flask populations (61–64). Competition between the two experimental ancestors revealed that the *ΔfimA* ancestor is extremely fit compared to the WT ancestor in both culture vessel environments (one-sample t-test, *P*_tube_ = 8.4 x 10-5, *P*_flask_ = 1.1 x 10-7, **Figure 3B**), particularly in tubes, where the WT ancestor was essentially undetectable after 24h of co-culture with the *ΔfimA* ancestor. However, competing the evolved WT populations against the *ΔfimA* ancestor revealed that WT populations surpassed the fitness of the *ΔfimA* ancestor by the end of the evolution experiment (one-sample t-test, *P*_tube_ = 0.047, *P*_flask_ = 0.0079) and exhibited a greater magnitude of fitness increase relative to the *ΔfimA* ancestor than the evolved *ΔfimA* populations. These results suggest that while the *ΔfimA* ancestor may have begun experimental evolution with an initial fitness advantage, there is still considerable room for fitness improvement and *ΔfimA-tube* populations may be stuck near a local fitness peak.

Finally, to determine if these evolved fitness outcomes were generalizable across environments, we assessed for signatures of local adaptation by repeating the competitive fitness assays in reciprocal environments (ex. assessed culture tube-evolved populations in flask environments). Contrastingly, all populations evolved from WT backgrounds exhibit patterns suggestive of local adaptation (ANOVA with Tukey’s HSD, *P*_Condition_ = 4.9 x 10^-4^, **Figures 3C-D**) while all *ΔfimA* populations exhibited greater fitness in flasks independent of their evolution environment. Thus, if competitive fitness in the tube environment is contingent upon the ability to form biofilms and to undergo range expansion, the perceived lack of increased fitness for *ΔfimA*-tube populations in their evolved spatially-structured environment may be indicative of a range specialist strategy that is revealed during culture in the flask environment which lacks spatial structure.

### Mutational parallelism illustrates genotype by environmental effects on evolutionary trajectories

Characterization of adaptive trajectories and determination of how genotype-by-environment interactions influence the effects of first-step mutations requires that we examine changes in the genome. To assess evolution on the molecular level and look for genetic patterns of positive selection, such as evidence of mutational parallelism, we performed metagenomic sequencing on the evolved populations at the 91-day timepoint (65). We also sequenced ancestor populations to remove any shared variation that could have been introduced when constructing the ancestral strains. After quality filtering to remove ancestral, low-quality, and low-frequency (<10%) polymorphisms, we identified 505 total mutations across all evolved populations (**Table 1, Table S1**). For all populations, nonsynonymous SNPs represented the largest proportion of mutations followed by intergenic SNPs, suggesting that the majority of mutations that increased to a frequency of > 10% resulted in an alteration of a gene’s function or expression as opposed to loss of function. One striking difference in the mutation spectrum was observed in mobile element insertions, where IS-elements are a greater contributor to evolution in tube populations compared to flask populations (**Figure 4A**).

**Table 1:**
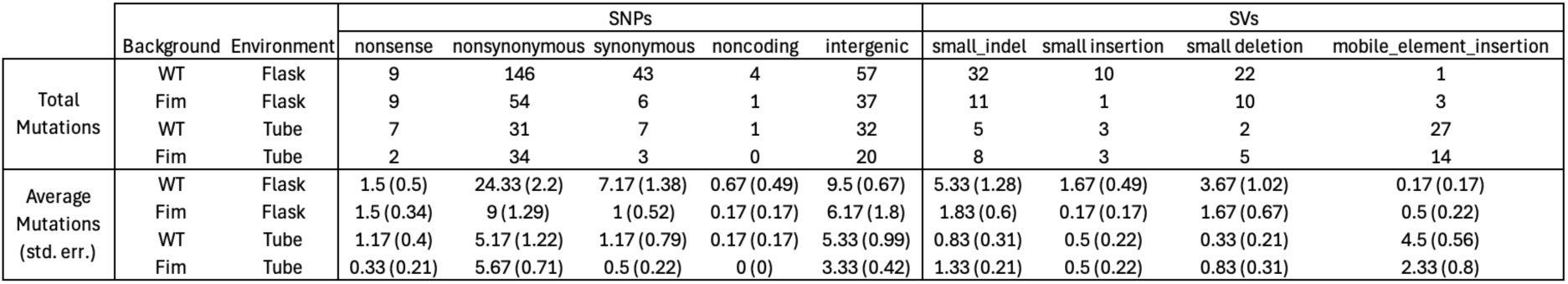
Mutations per treatment.

**Figure 4:**
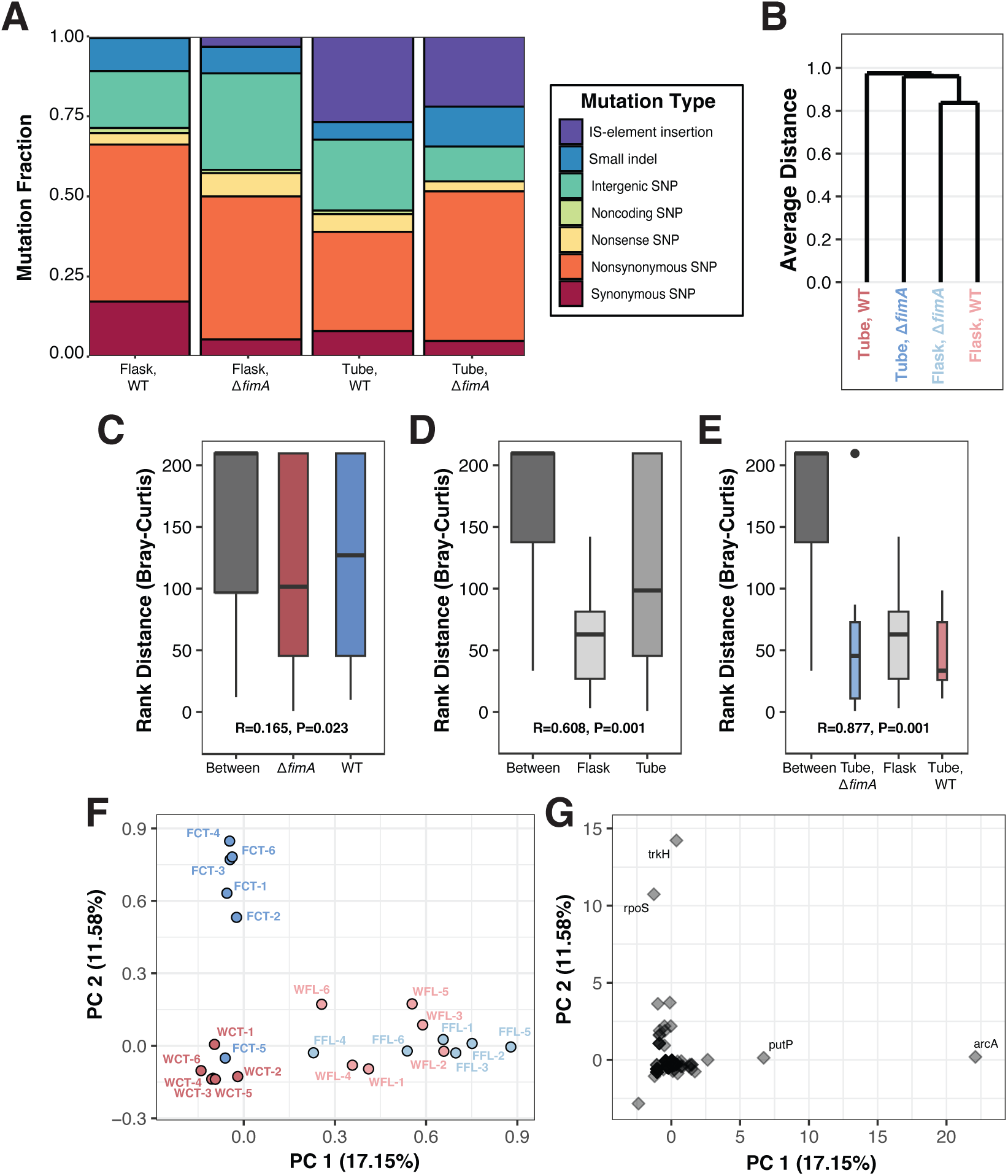
Mutational spectrum and parallelism are influenced by genotype-by-environment interactions. Metagenomic sequencing of evolved populations allowed the assessment of mutation spectrum and mutational parallelism. (A) Breakdown of mutations contributing to evolution in each genotype-environment combination grouped by mutation type. Colors indicate different mutation types. (B) Average mean mutational distance between each genotype-environment combination computed based on Bray Curtis dissimilarity. Rank distance based on Bray-Curtis dissimilarity comparing C) genetic background, D) evolution environment, and E) evolution environment with tube populations split by genetic background. For rank distance, lower values indicate greater similarity between grouped populations. F) Principal component plot of evolved populations and G) loci with nonsynonymous mutations. Principal component analysis was based on number of nonsynonymous mutations at each locus for each population. Panels F and G are from the same analysis and are plotted on the same coordinate space. For panels B - F, colors indicate each genotype-environment combination.

Considering these differences in the mutational spectrum, we also investigated gene-level mutational parallelism focusing on nonsynonymous mutations. To identify evidence of mutational parallelism, we first calculated average mean distance (**Figure 4B**) using the Bray-Curtis metric before conducting an analysis of similarity test (ANOSIM) to characterize the similarity of evolutionary trajectories within and between treatment groups (**Figures 4C-E**) (51, 66, 67). This analysis revealed that parallelism is best explained when flask populations are treated as a single group and *ΔfimA*-tube and WT-tube populations are each treated as their own groups (ANOSIM: R = 0.877, P = 0.001, **Figure 4E**), suggesting that *ΔfimA*-flask and WT-flask populations share similar gene-level mutational profiles, but *ΔfimA*-tube and WT-tube populations exhibit distinct profiles. To investigate this similarity further, we conducted a principal component analysis using the gene-level nonsynonymous mutation matrix and found a similar pattern where the individual flask populations cluster together while the tube populations cluster based on initial genotype (**Figure 4F**). This observed clustering of *ΔfimA*-flask is consistent with directional selection on these populations and our hypothesis that adaptation in the *ΔfimA*-flask populations could be following a range-specialist strategy.

Additionally, by plotting the genes we can also gain insight into the characteristic mutations of each evolutionary treatment (**Figure 4G**). For instance, flask populations are associated with nonsynonymous mutations in *arcA*, a member of the ArcAB two-component system that regulates the switch from aerobic to anaerobic respiration (68). WT-flask and *ΔfimA-*flask populations both contain 12 SNPs in *arcA*, with the majority of *arcA* mutations affecting the response regulator receiver domain (69, 70)(**Figure S4**). Alternatively, *ΔfimA-*tube populations are associated with nonsynonymous mutations in *trkH* with 7 SNPs in this gene that encodes a K^+^/H^+^ symporter (71). These SNPs primarily affect residues within the unresolved cytoplasmic loop of the TrkH protein (72). In addition to mutations in trkH, *ΔfimA-*tube populations also have 7 SNPs and a small insertion within *rpoS*. with five of these SNPs affecting residues that interface between the RpoS general stress sigma factor and the core RNA polymerase (73–76) (**Figure 5A**). Finally, while no true characteristic SNPs were identified via PCA for the WT-tube populations, two genes stood out in the genomic data as key targets of IS-element insertions, *fimE* (*fimA* regulating recombinase, IS-1 insertions) and *nlpD* (divisome associated factor, IS-5 insertions), consistent with the original evolution experiment that motivated this study (46, 50, 77, 78).

**Figure 5:**
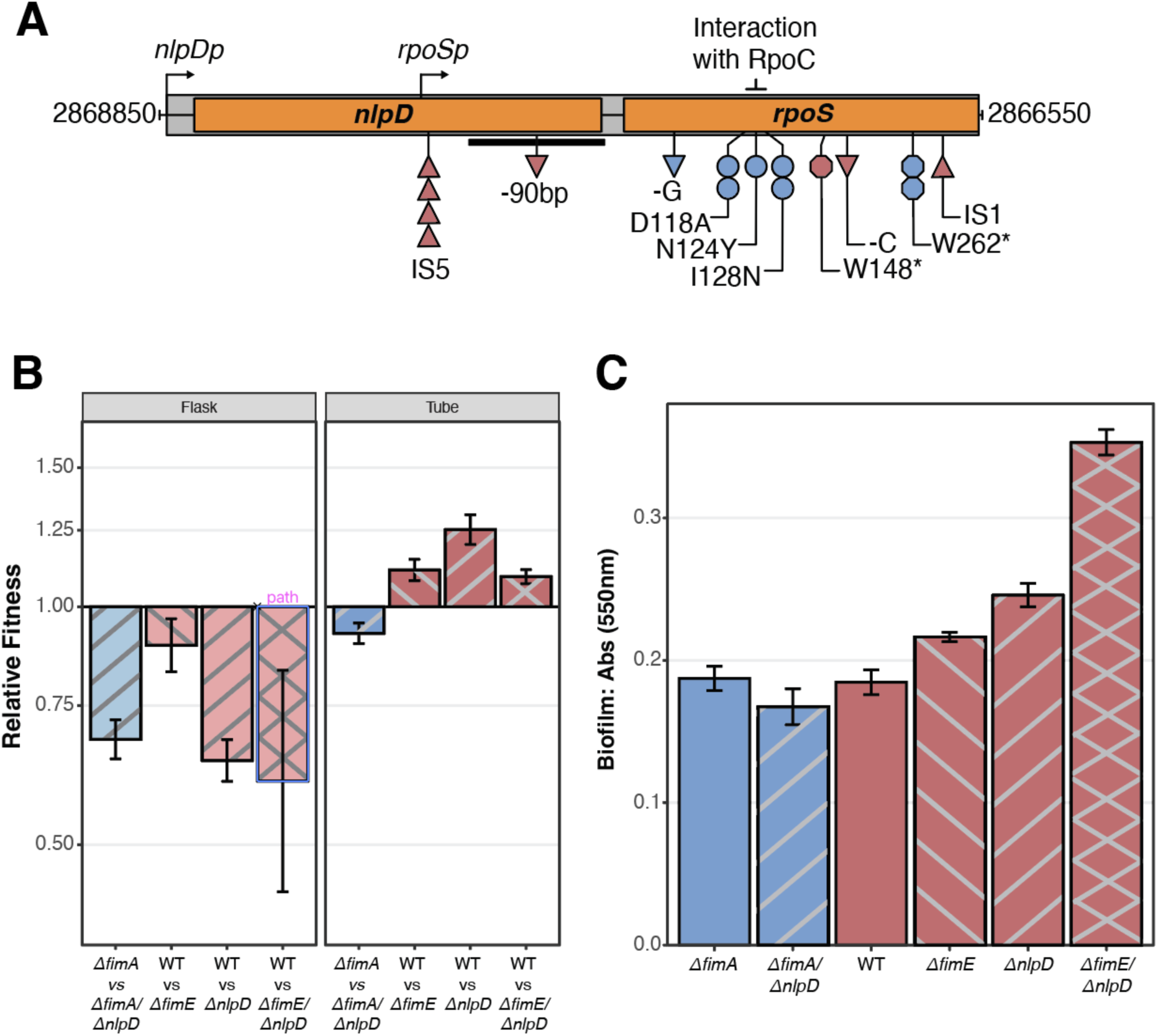
Mutations disrupting *nlpD* are beneficial in tube environments and contribute to biofilm formation. Fitness via competitive co-culture in the evolved environments and biofilm formation of *ΔnlpD* mutants was assessed. (A) Genic region of *nlpD* and *rpoS* loci shown, highlighting mutations observed in this experiment indicated below the labeled genes. Shapes indicate mutation types: insertions, upright triangles; deletions, upside-down triangles; SNPs, circles; nonsense mutations, octagons; colors indicate evolutionary treatment: red, WT-tube; blue, *ΔfimA*-tube. Notably, a *rpoS* promoter (*rpoSp*) embedded within the *nlpD* locus lies upstream of the insertion sites for the IS elements observed in our evolved populations. (B) Competitive co-cultures were conducted to assess the fitness impact of a *nlpD* deletion in combination with *ΔfimA* and *ΔfimE* mutations. After 24 hours of co-culture, results indicate that *nlpD* deletion confers a fitness advantage in the WT and *ΔfimE* background in the tube environment, but not in the flask environment. (C) Biofilm formation assays of *ΔnlpD* mutants, both alone and in combination with *ΔfimE* and *ΔfimA*, reveal an additive effect between the *ΔfimE* and *ΔnlpD* mutations. This finding helps to explain the recurrent emergence of these mutations in populations adapted to the tube environment. Competitive co-cultures and biofilm assays were performed at least 3 independent times at 37 °C. Error bars represent 95% confidence intervals of the mean.

### Insertions in *nlpD* are uniquely beneficial in WT backgrounds

As we observe the loss of FimE and NlpD function only in the WT-tube populations and not the *ΔfimA-*tube populations, these distinct mutations are key candidates for historically contingent adaptive trajectories in the tube environment. Interestingly, not only are *nlpD* and *rpoS* encoded in the same operon, but the coding region for *nlpD* also contains the *rpoS* promoter - located just upstream of the IS-5 insertions disrupting *nlpD* (77)(**Figure 5A**). As such, WT-tube populations may be taking a “two-birds-one-stone” approach as disruptions of *nlpD* will also disrupt *rpoS* expression (79). Thus, we expect that ruggedness in the fitness landscape may explain why *ΔfimA-*tube populations evolve mutations that directly disrupt *rpoS* while WT-tube populations instead evolve mutations in *nlpD* (*24, 80, 81*). To test this, we constructed *nlpD* and *fimE* deletions, individually and in tandem, in the ancestral WT background, as well as a *nlpD* deletion in the ancestral *ΔfimA* background. Competing these mutants against the WT and *ΔfimA* ancestors in the tube and flask environments revealed that deletions of *nlpD* were only beneficial in the tube environment, and only when present in the WT background or when combined with a *fimE* deletion (**Figure 5B).** We further assessed how these mutations contribute to biofilm formation and found that *ΔnlpD* significantly increases biofilm production when combined with *ΔfimE* in the WT background (**Figure 5C**)(pairwise t-test, *P_ΔfimE_* _vs. *ΔfimE/ΔnlpD*_ < 2.2 x10^-16^; *P_ΔnlpD_* _vs. *ΔfimE/ΔnlpD*_ = 6.4 x10^-12^), confirming the contribution of these mutations to enabling range expansion and highlighting how genotype-by-environment interactions influence evolutionary trajectories.

## Discussion

Here, we investigate the importance of genotype-by-environment interactions in determining the direction of evolution. As a first-step mutation, transposition of IS-1 into the *fimE* locus results in increased biofilm formation via the overexpression of the *fim* operon, enabling range expansion and colonization of the surface-air interface within the culture tube environment. However, an alternative first-step deletion of *fimA* is also initially beneficial as it reduces the energetic burden of baseline fimbriae expression. As such, we asked how first-step mutations can affect the adaptability of a population. By replaying evolution for 91 days, we observed that disruption of the *fim* operon is initially beneficial but ultimately deleterious and hinders adaptation to the structured environment by prohibiting range expansion. These effects are environmentally dependent, as deletion of *fimA* did not appear to affect adaptability in flasks, where biofilm formation is not advantageous, and WT populations ultimately evolve to repress biofilm formation. On the genomic level, we also observe genotype-by-environment effects on mutational parallelism as populations evolved in flasks exhibit more similar molecular evolution. In contrast, the molecular adaptive trajectories of tube populations are distinct between genetic backgrounds, as they both disrupt *rpoS* but through mutationally different means. Ultimately, our findings illustrate environmental effects on historical contingency and how initially beneficial first-step mutations can result in evolutionary traps that preclude future adaptation.

Prior research focused on first-step mutations have reported that these mutations often have large effects through adaptive loss of function (82, 83), widespread alteration of gene-regulation (84–86), and sign-epistasis (15, 87, 88). The effect of adaptive loss of function is most apparent in WT-tube populations where transpositions of IS-elements disrupting *fimE* and *nlpD* are the most common first and second-step mutations, sweeping to average frequencies of 0.813 and 0.706, respectively. These high frequency alleles are associated with biofilm formation, indicating that the highest population density is at the range margin, or surface-air interface biofilms, and that there is a greater initial fitness advantage associated with range expansion. As we previously observed, biofilm forming clones containing IS-insertions in *fimE* and/or *nlpD* regularly exhibit slower diauxic growth (50). Thus, it’s unlikely that this adaptive gene loss is associated with other cellular advantages, such as minimizing energetic waste.

Theory of r-K dynamics in range expansion predicts K-selection (higher density and slower growth rates) in the range core, and r-selection (lower density and higher growth rates) in the range margin (89). However, our study demonstrates the reverse to be true and other experimental evolution studies have similarly reported evidence of K-selection in range margins (90). For example, Fronhorfer and Altermatt observe high densities in *Tetrahymena population* range margins which they attribute to dispersal-foraging trade-offs as dispersal reduces competition (90). The inability to range expand essentially confines all individuals in *ΔfimA*-tube populations to the range core. Thus, the observation of *ΔfimA*-tube populations ultimately exhibiting higher fitness in flask environments is consistent with increased selection on foraging, as foraging efficiency and growth rate are expected to be greater contributors to competitive fitness in more homogeneous and unstructured conditions (91). This is further supported by the observation that genes with a role in resource uptake and utilization are enriched for higher frequency mutations in *ΔfimA*-tube populations, such as nonsynonymous mutations in *trkH* (K^+^:H^+^ symporter), and disruptive structural variants in the global regulators *cytR* (cytidine regulator) and *rpoS* (general stress response sigma factor). RpoS in particular has a major role in regulating growth and foraging as loss of *rpoS* decreases stress resistance but increases growth and resource utilization (92), while CytR represses genes involved in nucleoside uptake and scavenging (93). Interestingly, *nlpD*, which acquires IS-element insertions in all WT-tube populations, is located upstream of *rpoS* on the same operon and contains a dedicated *rpoS* promoter within its sequence (77, 79, 94). Thus, disruptions of *nlpD* serve a dual role as they contribute to biofilm formation (**Figure 5C**), but also downregulate expression of *rpoS,* resulting in broad-scale alterations of gene expression. Other studies have seen historical contingency result in divergent molecular but convergent phenotypic evolution (22, 95). As such, the divergent molecular but convergent phenotypic evolution of mutations disrupting *nlpD* in WT-tube populations versus mutations disrupting *rpoS* in *ΔfimA*-tube populations further illustrates how historical contingency influences the navigation of the genotype x phenotype landscape.

In addition to the fluctuating conditions used in our studies (50), mutations in *nlpD* and *fimE* were similarly observed during experimental evolution in tubes under constant 37°C temperature (51, 52). Here, out of 16 populations, all evolved mutations disrupting *fimE* and 8 evolved mutations disrupting *nlpD*. As these studies were independently conducted and initiated within a span of ∼5 years with slightly different conditions, this serves an excellent example of how selection can promote early parallel evolution. While this is especially true of mutations mediated by IS-element insertions and small indels, as these events are enriched within specific genomic locations, we also observe a number of mutations that are parallel on the nucleotide level.

Although mutational parallelism at the level of nucleotide position is thought to be rare, there are unique instances when we expect to observe this type of parallelism. 1) Stressful conditions with very strong selection: antibiotic stress (18), heat stress (96, 97), starvation stress (51, 98, 99), pH stress (100, 101). 2) Early polymorphic parallelism due to clonal interference (65). This is best exemplified in the LTEE where 312 mutations are detected in 2 or more populations at the same nucleotide position (129 nucleotide positions) within the first 2000 generations (**Table S2, Figure S5A**). Of these, only 4 mutations ultimately sweep to fixation, representing 1 nucleotide position (*pykF* A301S: Ara+1, Ara+5, Ara-5; *pykF* A301T: Ara-6). Similarly, in this study we observe 97 mutations that occur in 2 or more populations at the same nucleotide position (36 nucleotide positions) (**Figure S5B**). Of these, 54 mutations occur across multiple genotype x environment combinations (19 nucleotide positions) and only 1 has swept to fixation (*arcA*, F105L, WT-flask - population 1). As such, by sequencing early, we can detect early polymorphic parallelism before it is ultimately erased by the fixation of a singular allele.

Lastly, one surprising outcome of this study is that despite the evolutionary disadvantage, deletion of *fimA* is not rapidly compensated by expression of cryptic fimbriae in the tube environment. In *Escherichia coli*, there are eleven cryptic fimbriae that are similar to the type 1 fimbria encoded by the fim operon, but are not normally expressed by *E. coli* K-12 under laboratory conditions (45). Prior studies have demonstrated that at least six of these cryptic fimbriae encode functional adhesins when artificially expressed (45). In our study, *ΔfimA*-tube populations were unable to evolve expression of cryptic fimbriae which are typically repressed by H-NS, a pleiotropic histone-like nucleoid-binding protein and global repressor that regulates more than 5% of the *E. coli* genomes (60, 102, 103). As such, the ability to evolve expression of cryptic fimbriae may be under extreme genetic constraint as alteration of H-NS may result in severe unintended consequences of spurious gene expression or defects in chromosome organization. Because of this, deletion of *fimA* may be useful for studies using chemostats which are often complicated by biofilm formation (104), or FimA may prove to be a useful non-lethal target for therapeutic development (105). Ultimately, our study demonstrates how a seemingly beneficial first-step mutation can act as an evolutionary trap, stranding a population on a local adaptive peak.

## Materials and Methods

### Strain construction and experimental evolution

To follow up on evolved outcomes observed in Behringer et al., 2018 (50), we replayed evolution for 91 days (215 generations) to determine the effects of genetic background and environmental heterogeneity on *E. coli* adaptation. Here, we experimentally evolved a total of 24 parallel populations of *Escherichia coli* K-12 strain MG1655 that were evenly split across 4 different treatments: WT-tube, WT-flask, *ΔfimA*-tube, and *ΔfimA*-flask. All “WT” *E. coli* populations are descendants of PFM2, a prototrophic derivative of *E. coli* K-12 strain MG1655 (MG1655, *rph*+) (106). All *ΔfimA* cultures were constructed on the PFM2 background via P1 transduction by transferring the *fimA*782(del)::*kan* cassette from JW4277-1 (107) into PFM2 and disrupting the *fimA* gene. Additionally, to allow for the direct comparison of evolved fitness effects and easily screen for cross contamination, all WT-flask and *ΔfimA*-tube populations also contained a deletion of the *araBAD* operon (Δ*araBAD*567). Disruption of *araBAD* is a known neutral marker commonly used in microbial experimental evolution which allows strains to be colormetrically differentiated on Tetrazolium/Arabinose agar (TA) (108).

To conduct experimental evolution, transfers were conducted as previously described alternating 48h of growth at 25°C and 24h of growth at 37°C, shaking at 180 rpm (50)(**Figure 1**). Populations were each initiated by inoculating an independent isolated colony from its respective ancestral background into 10 mL of LB broth. Each subsequent transfer consisted of a 1:10 dilution where 1 mL of well-mixed evolving culture was inoculated into 9 mL of fresh LB broth, resulting in ∼ 3.3 generations per transfer. Populations evolving to culture tube treatments were cultivated in 16 x 100 mm glass culture tubes (VWR, Cat. 47729-576) while populations evolving to flask treatments were cultivated in 50 mL glass culture Erlenmeyer flasks (Pyrex, Cat. 4980). Every week following a 24h/37°C incubation, populations were screened for contamination by streaking 1 μL of culture on TA and McConkey agar plates. To maintain a rich historical record of evolution, each week following the succeeding 48h/25°C incubation 1 mL of well-mixed pre-transfer culture was collected and stored in 40% glycerol at -80°C. Following the 91-day time point, additional 1 mL aliquots were collected for DNA isolation and metagenomic sequencing. Here, the spent media was removed via centrifugation (10,000 x g for 2 m) and the culture was flash frozen with liquid nitrogen before storing at -80°C.

To assess phenotypes of mutations identified in this work, we generated *ΔnlpD* and *ΔfimE* mutants in multiple genetic backgrounds. The *ΔnlpD* single mutant strain was generated by moving the *nlpD*747(del)::*kan* cassette from the Keio collection strain JW2712 to PFM2 (WT) using P1 transduction. Similarly, the *ΔfimE* single mutant strain was created by moving the *fimE*781(del)::*kan* cassette from the Keio collection strain JW4276 to PFM2 (WT) using P1 transduction. To produce the *ΔfimE/ΔnlpD* and *ΔfimA/ΔnlpD* double mutant strains, the kanamycin resistance cassette was first excised from the *ΔfimE* and *ΔfimA* single mutant strains. This was accomplished by expressing the FLP recombinase from the helper plasmid pCP20, which resulted in the removal of the kanamycin resistance cassette leaving behind the FRT scar site (109).

Subsequently, the *nlpD*747(del)::*kan* cassette from JW2712 was introduced into the resulting *ΔfimE* and *ΔfimA* strains through P1 transduction, resulting in the *ΔfimE/ΔnlpD* and *ΔfimA/ΔnlpD* strains. All mutants were verified via PCR followed by Sanger sequencing.

### Measurement of Growth Phenotypes and Quantification of Biofilm Formation

Pre-transfer cell density was measured weekly before a 24h/37°C transfer and a 48h/25°C transfer over the course of experimental evolution by adding 150 μL of well-mixed culture to a 96-well plate (Corning/Falcon, Cat. 351172) and then recording absorbance at 600 nm in an Epoch2 Microplate Spectrophotometer (Biotek). Cell density measurements for each evolving population were taken in duplicate at each timepoint. To quantify evolved differences in the ability to form biofilms, we used a microtiter plate biofilm assay (59). For biofilm formation at 25°C, 15 μL of 25°C/48h culture was inoculated into 96-well plates (Corning/Falcon, Cat. 351172) containing 150 μL of LB broth. Inoculated 96-well plates were incubated for 48 h at 25°C before removing the spent media containing unattached cells and washing the plate three times with 1x PBS. Plates were then stained with 200 μL of 0.1% crystal violet for 10 m before washing again with 1x PBS and left to dry overnight. Crystal violet bound to biofilms was then solubilized with 200 μL 30% acetate for 15 m and transferred to a fresh 96-well plate. Absorbance was quantified using an Epoch2 Microplate Spectrophotometer at 550 nm. To compare biofilm formation between 48h/25°C treatments and 37°C/24h treatments, this process was repeated with incubation temperatures adjusted to 37°C and incubation times adjusted to 24h. Each evolved population/timepoint combination was measured for a total of 8 replicates and each 96-well plate assay contained 8 replicates of the WT ancestor (PFM2) to serve as a control. To determine if differences in biofilm formation could be attributed to differences in cell density, biofilm measurements were normalized to cell density.

### Estimation of Relative Fitness

To determine how biofilm formation affects adaptation to different environments, we assessed relative fitness for each 91-day evolved population in comparison to its ancestor (57). Fitness was measured in both the evolved and the reciprocal environments to assess the degree of local adaptation. 48 hours before the start of the competitions, the ancestor strains were streaked out on LB agar from the frozen stock and incubated at 37℃ for 24 hours. 24 hours before the start of the competitions, overnight monocultures were made in the competition environments for the evolved strains from the frozen stock and for the ancestor strains from the plate cultures. The overnight cultures were incubated at 37℃ for 24 hours. The optical density of these cultures was measured using an Epoch2 Microplate Spectrophotometer at 600 nm and used to calculate the cell density in the overnight cultures (**Equation 1**) to equalize the concentration of evolved and ancestor cells in 10 mL LB broth.

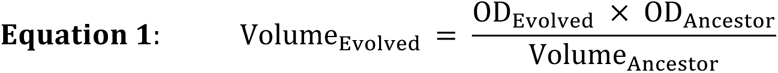

The tubes were then vortexed using a sterile protocol, and the flasks were mixed by swirling. 100 μL from each competition were transferred to a microcentrifuge tube with 900 μL 1x PBS, and dilutions from these samples were done up to a factor of 10^-5^. The co-culture competition tubes and flasks were incubated at 37℃ and 25 ℃ for 24 h and 48 h, respectively. The 10^-4^ and 10^-5^ dilutions were each plated on TA agar using 100 μL from each microcentrifuge tube. After 24 hours, the plate counts (differentiated by an *araBAD* disruption as detailed above) were used to determine the colony forming units (CFU) of each cell type in the competition at 0 hours (**Equation 2**). Final plate counts were determined in the same manner, except that dilutions were done up to a factor of 10^-6^.

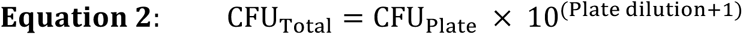

The initial and final cell counts for each competition were used to determine the relative fitness (*W*) of each strain (110) (**Equation 3**).

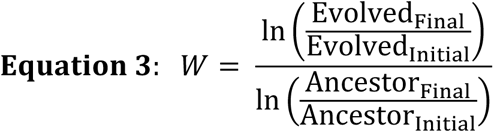

Determining the relative fitness of each evolved strain when compared to its ancestor shows the extent to which 91 days of experimental evolution in a specific environment affects fitness in that environment. However, we also competed WT populations against *ΔfimA* populations in both environments to determine how genetic background affects fitness. The same protocol was used as detailed above, but these competitions occurred between ancestors or between evolved populations.

### DNA Extraction and Mutation Calling

DNA was extracted from the previously frozen copies of each line using the DNeasy UltraClean Microbial Kit (Qiagen). Library preparation and metagenomic sequencing were performed at the Vanderbilt University Medical Center. The samples were normalized to target 100 ng of total input for each sample, and libraries were prepared using the Twist Biosciences kit (Twist, 104207) following a miniaturized version of the manufacturer’s protocol. The libraries were quantitated and pooled for downstream processing. The pool quality is assessed using the Agilent Bioanalyzer and quantified using a qPCR-based method with the KAPA Library Quantification Kit (Kapa, KK4873) and the QuantStudio 12K instrument. Prepared libraries were pooled in equimolar ratios, and the resulting pool was subjected to cluster generation using the NovaSeq 6000 System, following the manufacturer’s protocols. 150 bp paired-end sequencing was performed on the NovaSeq 6000 platform targeting 2M reads per library. Raw sequencing data (FASTQ files) obtained from the NovaSeq 6000 was subjected to quality control analysis, including read quality assessment. Real Time Analysis Software (RTA) and NovaSeq Control Software (NCS) (1.8.0; Illumina) were used for base calling. MultiQC (v1.7; Illumina) was used for data quality assessments.

Resulting reads were processed using fastp v.0.23.1 to remove residual adapters and trim low-quality sequences (111). After this quality control process, we mapped population-level metagenomic sequencing reads to the reference genome of *Escherichia coli* K-12 substr. MG1655 (NC_000913.3) with the Breseq v.0.35.0 pipeline, calling mutations and their frequencies with the predict-polymorphisms parameter setting (112). Before statistical analysis, mutations present in each ancestor population were filtered out of the corresponding evolved populations, as well as low quality and low-frequency polymorphisms (<10%). We chose 10% as this was the cutoff where we minimize calling artifacts from library preparation.

### Statistical Analysis of Genomic Data

To identify parallel evolution within and among evolutionary treatments, we focused on nonsynonymous SNPs. Here, we constructed a matrix of every gene containing a non-synonymous SNP in at least one population and counted the number of SNPs in that gene for each population. Using this matrix we assessed mean genetic distance and conducted an analysis of similarity within and among treatments using the Bray-Curtis method with the vegan package in R (113). Using the same matrix, we then performed principle component analysis to assess between population similarity and identify characteristic mutations associated with each treatment using the PCA function from the FactoMineR package in R (114).

## Acknowledgements

We would like to thank M.A., R.F., M.L., L.G., and A.U. for their scientific insight and contributions to this study. This work was supported by National Science Foundation grant #2117782 awarded to M.G.B.; start-up funds from the Vanderbilt University College of Arts and Science and the Vanderbilt Institute for Infection, Immunology and Inflammation; as well as a Vanderbilt Undergraduate Summer Research Fellowship sponsored by the Littlejohn Family awarded to (G.F.P.). We declare no conflicts of interest.

## Author Contributions

M.G.B. conceived and designed the study. M.G.B., R.S., and S.B.W. constructed the strains. M.B. and J.C.M. performed experimental evolution and collected cell density data. M.G.B., S.B.W. and S.W. assessed biofilm formation. M.G.B., G.P., and M.Y. assessed competitive fitness. M.G.B. and G.P. conducted the genomic analyses. M.G.B. and G.P. wrote the manuscript.

## Supplemental Figures and Tables

**Figure S1:**
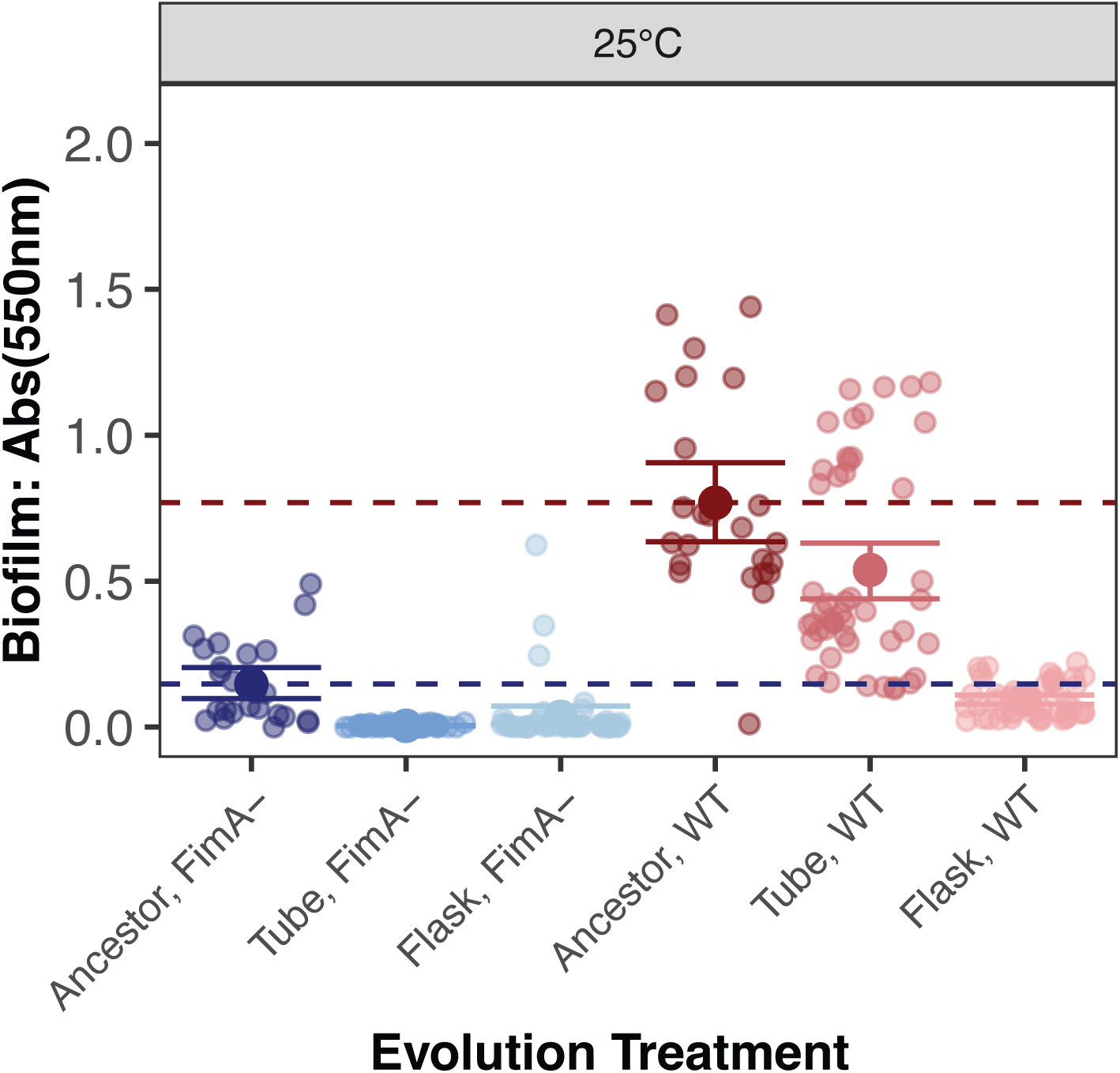
Biofilm production after 48h culture at 25°C. Red dashed line indicates the mean value for the WT ancestor, blue dashed line indicates the mean value for the *ΔfimA* ancestor. Error bars represent 95% confidence interval of the mean.

**Figure S2:**
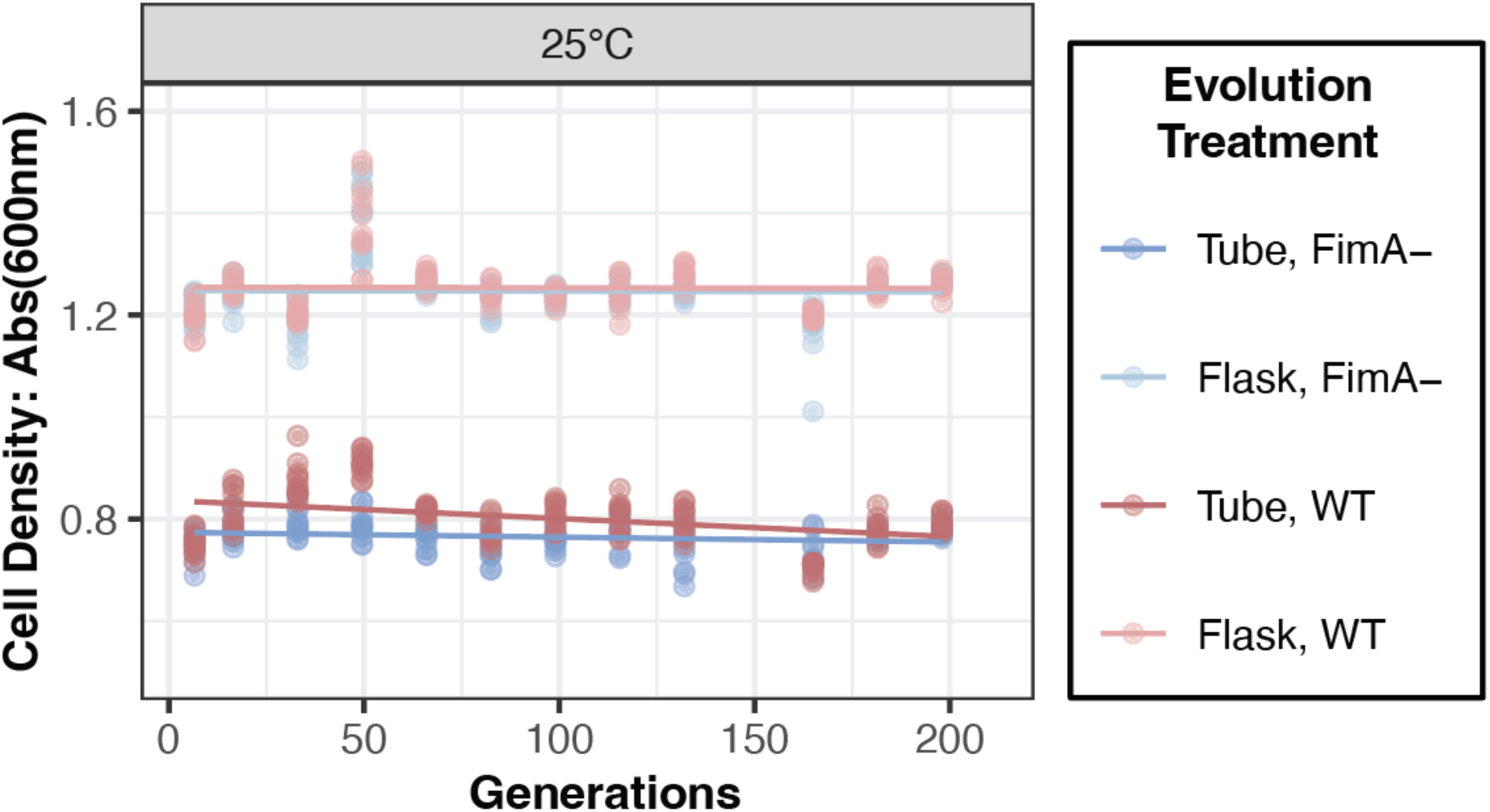
Pre-transfer culture density throughout experimental evolution after 48h culture at 25°C. Lines indicate a linear fit of the change in density over time.

**Figure S3:**
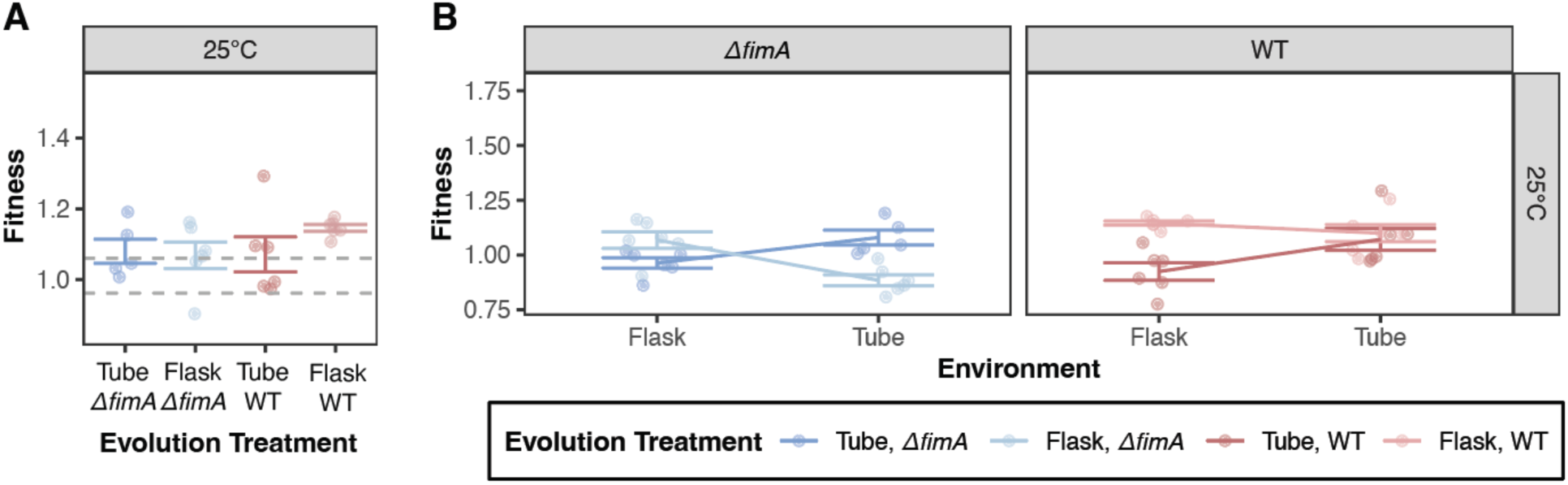
Fitness assessment following 48h co-culture at 25°C. Fitness is assessed by competitive co-culture in evolved and reciprocal environments. (A) Fitness of each population in their evolved environment after 91 days relative to the ancestor strain of the same genetic background (e.g. evolved *ΔfimA*-tube with *ΔfimA* ancestor). (B) Fitness of each population in their evolved or reciprocal environment relative to the ancestor strain of the same genetic background. Lines illustrate fitness relationships by connecting the mean fitness for each evolved genotype-environment combination, error bars represent 95% confidence intervals of the mean.

**Figure S4:**
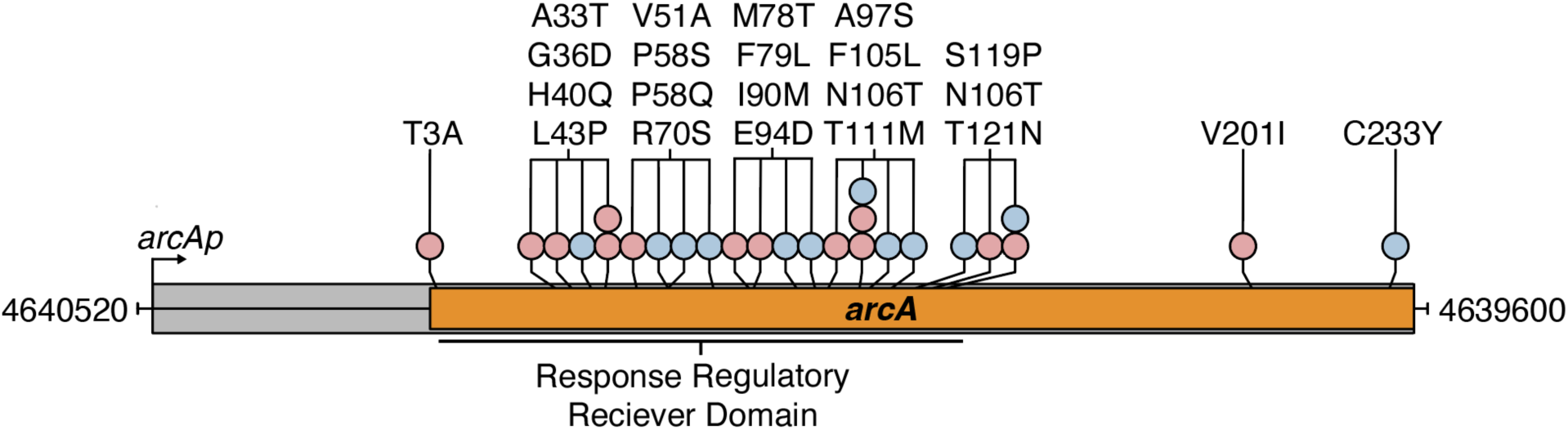
Genomic location *arcA* mutations enriched in flask populations. Genic region of *arcA* locus shown, highlighting mutations observed in this experiment indicated above the labeled genes. Shapes indicate mutation types: SNPs, circles; colors indicate evolutionary treatment: red, WT-flask; blue, *ΔfimA*-flask.

**Figure S5:**
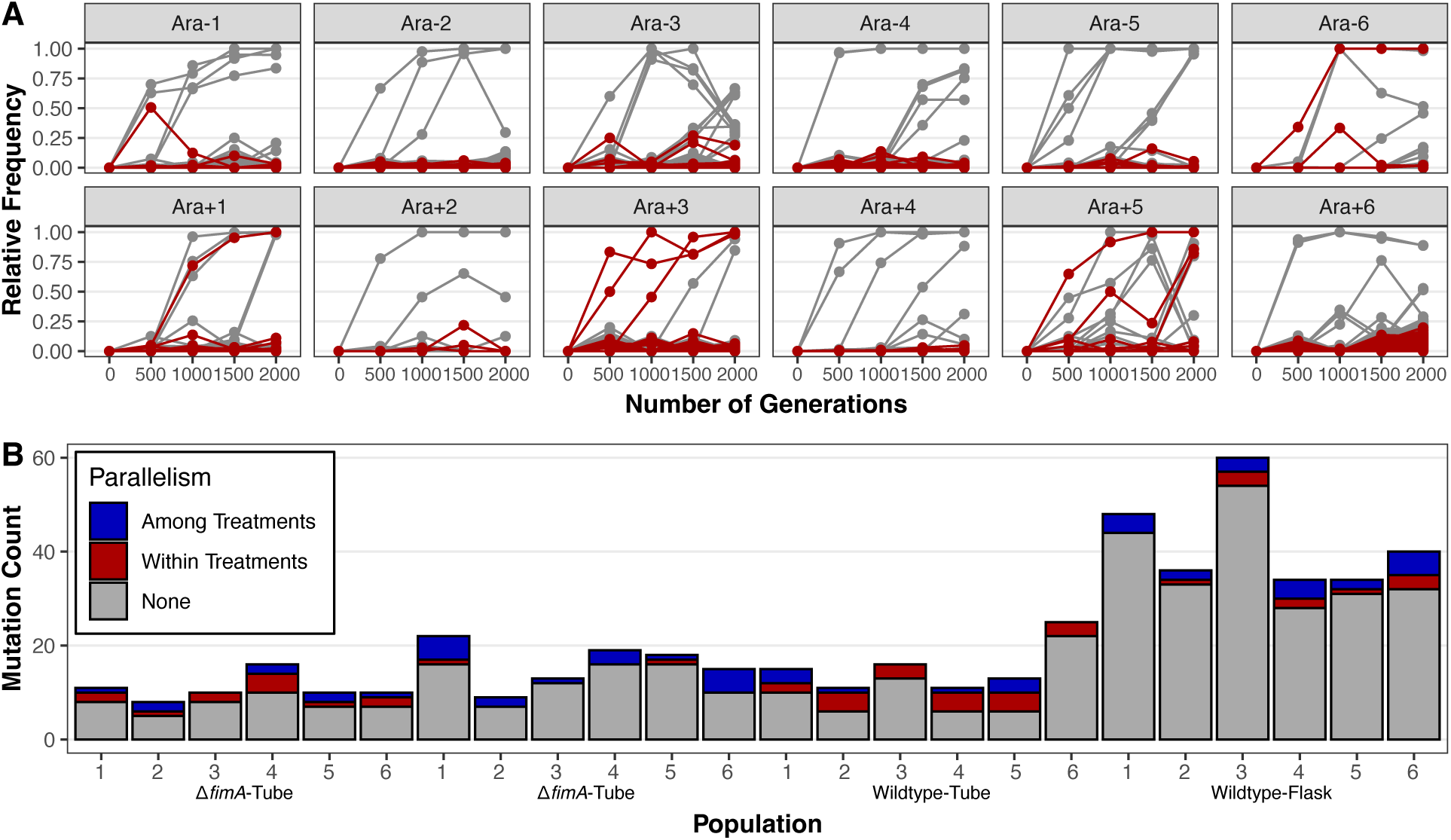
Early nucleotide-level parallelism in experimental evolution datasets. A) Parallel polymorphisms across the first 2000 generations of evolution in the LTEE. Gray points and lines indicate unique polymorphisms, red points and lines indicate polymorphisms shared across two or more populations. B) Parallel polymorphisms in this experiment. Grey bars indicate unique polymorphisms, red bars indicate polymorphisms shared within an evolution treatment, blue bars indicate polymorphisms shared across multiple evolution treatments.

Table S1: All mutations over 10% frequency for each population

Table S2: Early parallelism in the LTEE

